# Forensic analysis of Soil Microbiomes: Linking Evidence to a Geographic Location

**DOI:** 10.1101/2020.07.10.198044

**Authors:** Jill Hager Cocking, Sgt. Ryan Turley, Viacheslav Y. Fofanov, Kimberly Samuels-Crow, Bruce Hungate, Rebecca L. Mau, Paul S. Keim, J. Gregory Caporaso, Crystal Hepp

## Abstract

Over the past two decades, advances in molecular biology have greatly expanded our understanding of microbiomes – the diverse assemblages of microorganisms that inhabit the human body as well as the world around us, and applications in microbiome science have become an active area of research. Differences in the diversity (i.e., richness) and composition of microbiomes has been found to be informative in varied areas of science, including human health, agronomy, and forensic science. Soil harbors microbiomes that vary based on many factors, including the geology of the soil (e.g., sand, silt, or clay), climate, and use of the soil. As a result, the microbiological composition of any two soil samples will never be exactly alike. This inherent variation between microbiomes of different locations has proven to be specific enough to be potentially useful in forensic investigations to associate a person or piece of evidence to a source site.

In this study, a soil microbiome was extracted from the sock of a criminal suspect and compared to the microbiome of soil samples taken from locations traveled to by the suspect. The locations analyzed varied in their soil microbiome composition, and the microbiome profiled from the sock was found to be most similar to the location where the suspect was thought to have left the body of a murder victim. These results provide a case study illustrating that information contained in a soil microbiome may be applied to link evidence to the location where a crime took place, potentially serving as an investigative tool in law enforcement.

---

Recent advances in DNA sequencing and bioinformatics analysis methods have led to a better understanding of microbiomes, communities of different species of microscopic organisms whose metabolisms are tightly linked to one another, to their environments, or to their plant and animal hosts. These technological advances have led to our recent recognition that there are orders of magnitude more microbial species than were previously thought to have existed (1). This has led to a revolution in microbiology, with new sub-fields of science forming to understand the composition and functional activities of microbiomes. The microbiome of the skin and inside the human body varies substantially depending on the body location sampled and has revealed itself as an important indicator of health (2). For example, individuals with chronic conditions like Crohn’s Disease and chronic rhinosinusitis harbor gut (3) and sinus microbiomes (4), respectively, with lower richness (i.e., fewer different species present) than the microbiomes of healthy individuals. Similarly, individuals with acute viral gastroenteritis also have less diverse gut microbiomes (5). Although fecal supplementation following antibiotic treatment has seen fringe use in medicine since the 1950s (6), we are now beginning to formally develop applications of this knowledge in human health. For example, the transplantation of a healthy individual’s gut microbiome into a patient suffering from recurring *Clostridium difficile* infections is now becoming common (7).

Microbiome analysis techniques extend beyond human health and already have some recognized forensic applications. For example, skin microbiomes differ between human subjects, and enough of this unique microbial fingerprint is left behind on the objects that we touch that objects, such as keyboards and computer mice, can be linked to their owner solely based on the microbes found on their surfaces (8). Similarly, the bacterial community microbiome found in human saliva was shown to be unique to an individual (9). Even time of death information may be gleaned from the succession of microbial communities which are involved in decomposition (10). Taken together, these studies suggest a role for microbiome science in forensics applications.

Microbes, and bacteria and fungi in particular, are integral components of soil. The specific microbial composition of a soil sample is driven by factors including the physicochemical properties of the soil (e.g., pH, salinity) (11, 12), environmental features (13), and land use (14), such that no two soil samples will ever be exactly alike. This variation between soils from different locations has proven to be specific enough to be potentially useful in forensic investigations (15).

We therefore hypothesized that soil extracted from a piece of evidence may provide sufficient information to link that evidence to the soil’s source.

In October of 2017, our laboratory was contacted by the Flagstaff Police Department. A woman was missing and presumed dead. A suspect was in custody but was not revealing the location of the missing woman. Using various investigative tools, the police knew where the suspect had traveled since being released on bail from the Flagstaff jail a few days earlier. The police were trying to decide whether to focus their search efforts in the town of Mayer, Arizona (Yavapai County) or Williams, Arizona (Coconino County), with a distance between the two of 87 to 118 miles depending on the route taken. A sock embedded with soil, believed to have been worn by the suspect while not wearing a shoe, was in the custody of the police. Their hope was that the sock could be analyzed and linked to one of the two locations to aid in finding the body. During the following few days (before the microbiome analysis was completed), the body was recovered in Mayer, Arizona. Even though the police no longer needed assistance with the recovery of the body, we attempted to analyze soil embedded in the sock in order to compare it to various locations around Yavapai and Coconino Counties in the state of Arizona to determine if the soil microbiome could be a useful forensic tool in future investigations. Knowing that there are inherent variations in soil microbial community depending on location, we compared the bacterial community found on the sock with bacterial communities found in multiple soil samples collected from locations traveled to by the suspect during the days prior to his arrest.

## Methods

Reference surface soil samples were obtained from 18 locations around Williams (Coconino County) and Mayer (Yavapai Country), Arizona (Table 1). Locations were selected because they were either near the site where the body was found (Fig. 1) or were places the suspect was known to have traveled in the days after he was last seen with the victim. Five soil samples collected from other locations in Arizona were also obtained from the Center for Ecosystem Science and Society (Ecoss) at Northern Arizona University to serve as additional reference samples (the “Ecoss reference samples”). Together this resulted in 23 reference samples where K1-11 and K13 refer to the Mayer soil samples, K12, K14-16, and K18 refer to the Williams soil samples, K17 refers to the Chino Valley soil sample and W0.GL.1, W0.PJ.1, W0.MC.1, W0. PP.1 and 217 refer to the Ecoss reference samples.

**TABLE 1.** Information for samples used for analysis.

| Sample Name | Description |
| --- | --- |
| Q1 | small cutting from sock, ball of foot area |
| Q2 | small cutting from sock, heel area |
| Q3 | swabbing of visibly dirt-covered area of sock for 3 minutes |
| Q4 | swabbing of visibly dirt-covered area of sock for 3 minutes |
| K1 | soil, right off highway Mayer, AZ |
| K2 | soil, Road by trailer of owner of land where body was found, Mayer, AZ |
| K3 | soil, off Road A, Mayer, AZ |
| K4 | soil, dry, cracked ground near where body was found, Mayer, AZ |
| K5 | soil, near body site, Mayer, AZ |
| K6 | soil, 5 ft west of body site decomposition area, Mayer, AZ |
| K7 | soil, 5 ft north of body site decomposition area, Mayer, AZ |
| K8 | soil, 5 ft east of body site decomposition area, Mayer, AZ |
| K9 | soil, body site with visual decomposition residue, Mayer, AZ |
| K10 | soil, body site with visual decomposition residue, Mayer, AZ |
| K11 | soil, behind body site, more plants, Mayer, AZ |
| K12 | soil, Williams, AZ |
| K13 | soil, 5 ft south of body site decomposition area, Mayer, AZ |
| K14 | soil, Williams, AZ |
| K15 | soil, off Forest Service Road, Williams, AZ |
| K16 | soil, Forest Service Road, Williams, AZ |
| K17 | Soil, Chino Valley, AZ |
| K18 | Soil, Williams, AZ |
| W0.GL.1 | Grassland site of C. Hart Merriam Gradient, Northern Arizona |
| W0.PJ.1 | Pinyon-Juniper site of C. Hart Merriam Gradient, Northern Arizona |
| W0.MC.1 | Mixed Conifer site of C. Hart Merriam Gradient, Northern Arizona |
| W0.PP.1 | Ponderosa Pine site of C. Hart Merriam Gradient, Northern Arizona |
| 217 | Mixed Conifer site of C. Hart Merriam Gradient, Northern Arizona |
| Reagent Blank Sock (RBsock) | Reagent Blank extracted alongside sock samples |
| Reagent Blank Soil (RBsoil) | Reagent Blank extracted alongside soil samples |

**FIG. 1.**
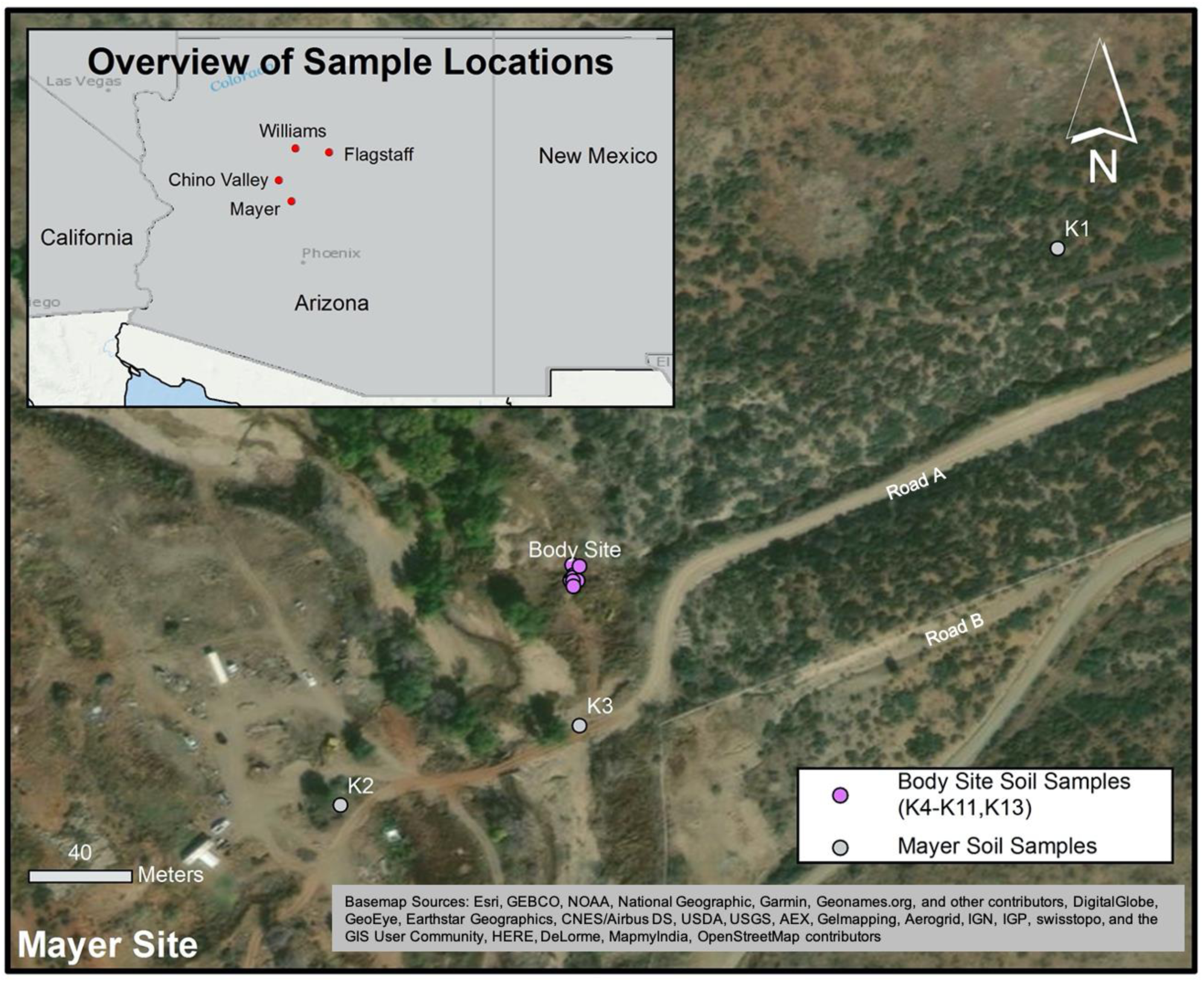
Locations of reference soil samples collected at and near the body site in Mayer, Arizona. Inset map shows geographic relationships between the Mayer site and other locations relevant to the case. Mayer Soil Samples include K1, K2, and K3. Body site soil samples include K4-K11 and K13 (Table 1).

The sock belonging to the suspect was visibly soiled over most of its surface. DNA extraction from the sock was performed in two ways with two replicates per extraction approach, yielding four query samples, referred to here as Q 1-4. Two DNA samples were extracted from cuttings of the sock itself (approximately 2 cm^2^ each). One cutting was taken from the ball of the foot area (Q1) and one cutting was taken from the heel area (Q2). For the second extraction method, 2 pre-moistened cotton swabs (Q3, Q4) were rubbed over the visibly soiled area of the outside of the sock for 3 minutes each, as described by Goga (16). The swabs were then removed from the applicator and used for the remainder of the extractions. The extractions were performed using the Qiagen DNeasy® PowerSoil Kit (Qiagen, Germantown, MD) according to the manufacturer’s protocol, with the following variations. The cuttings and swabs were placed at 65°C for 10 minutes followed by 2 minutes of horizontal vortexing at the maximum speed of the vortexer. The final elution volume was 100 µL.

For the reference soil samples (K1-18), approximately 0.25 grams of each of the 18 soil samples (Table 1) were added to a PowerBead tube containing solution C1. This tube was placed at 65°C for 10 minutes followed by 2 minutes of horizontal vortexing at maximum speed. The remainder of the extraction was performed according to the Qiagen DNeasy® PowerSoil Kit’s manufacturer’s instructions.

The EcoSS reference soil samples differed from the reference samples collected for this study in that they were collected below the soil surface (0-10 cm) while K1-18 were taken from the surface (as the surface soil would be the most likely to come into contact with the suspect’s sock). There is known variation in soil microbiome composition depending on sampling depth (17), but these samples were included to provide additional background soils to which we could compare our query samples.

Extractions from Q1-4 and K1-18 were performed at different times in 2017. DNA was extracted from the EcoSS reference samples in 2014 and 2015 by using a MO BIO PowerSoil(tm) DNA Isolation Kit (Qiagen, Germantown, MD) and following the manufacturer’s directions. Briefly, approximately 0.25 g of soil was added to the lysis tube and lysed using a MP Biomedicals FastPrep Homogenizer (MP Biomedicals, Irvine, CA). The final elution volume was 100 ul.

During the extractions for Q1-4 and K1-18, a reagent blank was taken through the entire process to monitor laboratory and extraction reagent contamination. For the sock samples, the reagent blank (RBsock) consisted of a cotton tipped swab moistened with UltraPure distilled H2O and cut with scissors used for the sock and handled with tweezers used for the sock extraction. The reagent blank was processed alongside the sock extractions. For the reference soil reagent blank (RBsoil), water was added to a weigh boat and then placed in a PowerBead tube and processed alongside the known soil samples.

The hypervariable V4 region of the 16S rRNA gene was amplified from each of the reference soil samples and sock samples, as well as the reagent blanks. This amplified DNA was then prepared for sequencing on the Illumina MiSeq instrument according to the protocol presented by Caporaso *et al* (18). The resulting sequences were analyzed using QIIME 2 microbiome bioinformatics platform (19). Sequence quality control was performed using the denoise-paired method of QIIME 2’s DADA2 (20) plugin with the following parameter settings: trunc_len_f 293; trunc_len_r 208; trim_left_f 6, trim_left_r 6. The resulting amplicon sequence variants were assigned taxonomy using q2-feature-classifier’s classify-sklearn method against (21) GreenGenes (22) 13_8. ASV sequences were aligned using MAFFT (23) (qiime alignment mafft), highly variable positions were filtered (qiime alignment mask), an unrooted tree was constructed using FastTree (24) (qiime phylogeny fasttree), and the tree was rooted by midpoint rooting (qiime phylogeny midpoint-root). Weighted and unweighted UniFrac (25) distances were computed 100 times each at an even sampling depth of 1000 sequences per sample. This low depth of coverage was used to retain all samples in the analysis, and 100 iterations were run to confirm that conclusions were robust across rarefied feature tables. These analyses were performed using the beta-rarefaction visualizer in QIIME 2’s diversity plugin. Sample tree illustrations were generated with ete3 (26).

## Results

DNA was successfully extracted and the V4 region of the 16S rRNA gene was amplified from the 4 sock samples, the 2 reagent blanks, and the 23 reference soil samples. Weighted and unweighted UniFrac neighbor joining trees were constructed to evaluate the similarity of microbiomes (Fig. 2a and 2b, respectively). Briefly, the UniFrac metrics provide distances between pairs of microbiome samples. Smaller values indicate that a pair of samples are similar in their composition, while larger values indicate that a pair of samples are dissimilar in their composition. The unweighted UniFrac metric is considered a qualitative metric in that it only compares samples based on which microbes are present, but does not consider the abundance of those microbes. The weighted UniFrac metric is considered a quantitative metric because it compares the abundances of different microbes in the samples. Because estimation of microbial abundances is imperfect using the techniques applied for microbiome profiling, both weighted and unweighted UniFrac metrics are often computed and compared. These metrics are applied to compute distances between all pairs of microbiome samples, and the resulting distance matrix can be summarized by constructing a neighbor joining tree. In this tree, samples are represented as the leaves (or tips), and the length of the branches between leaves represents the distance between the samples.

**FIG. 2.**
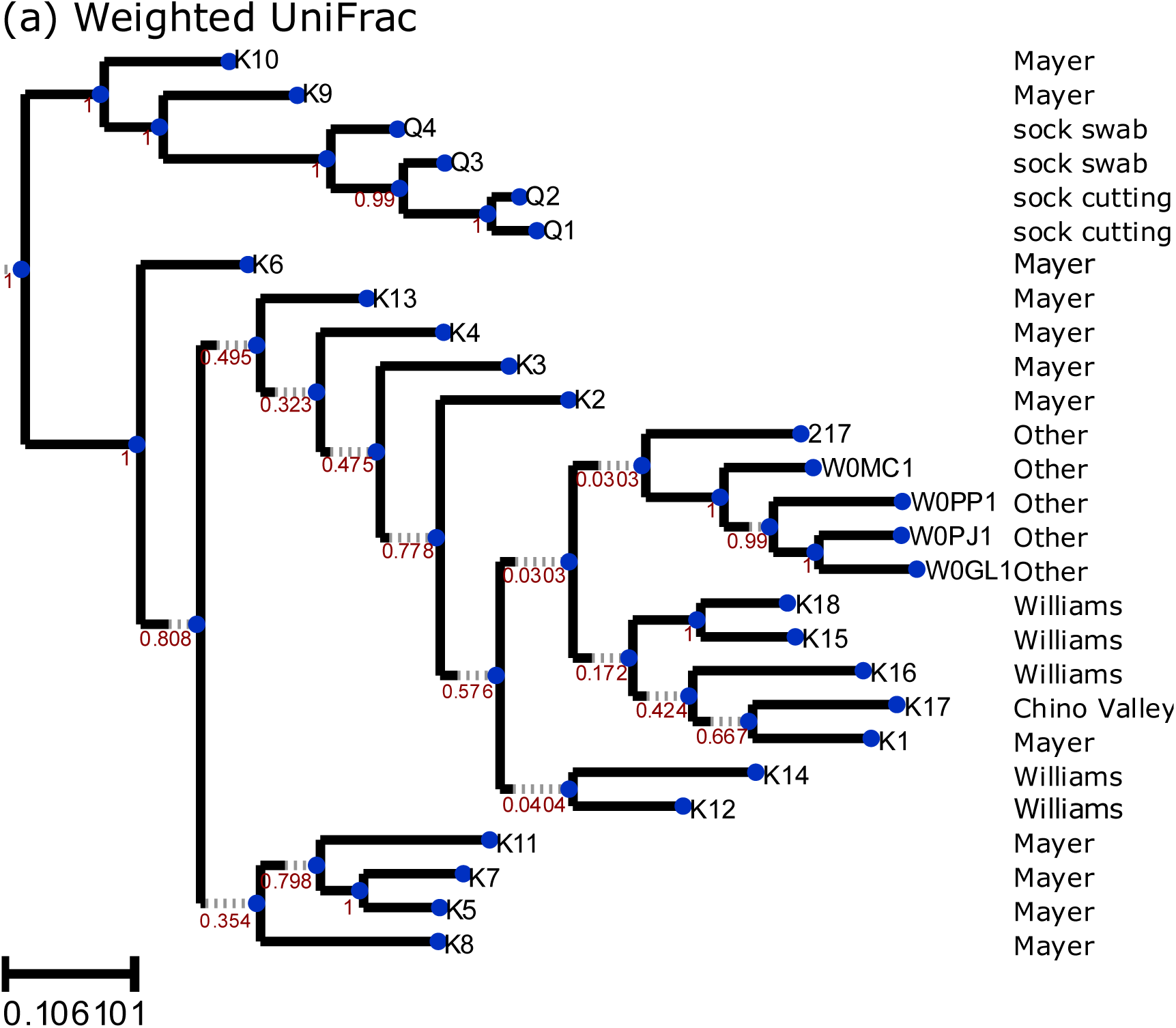
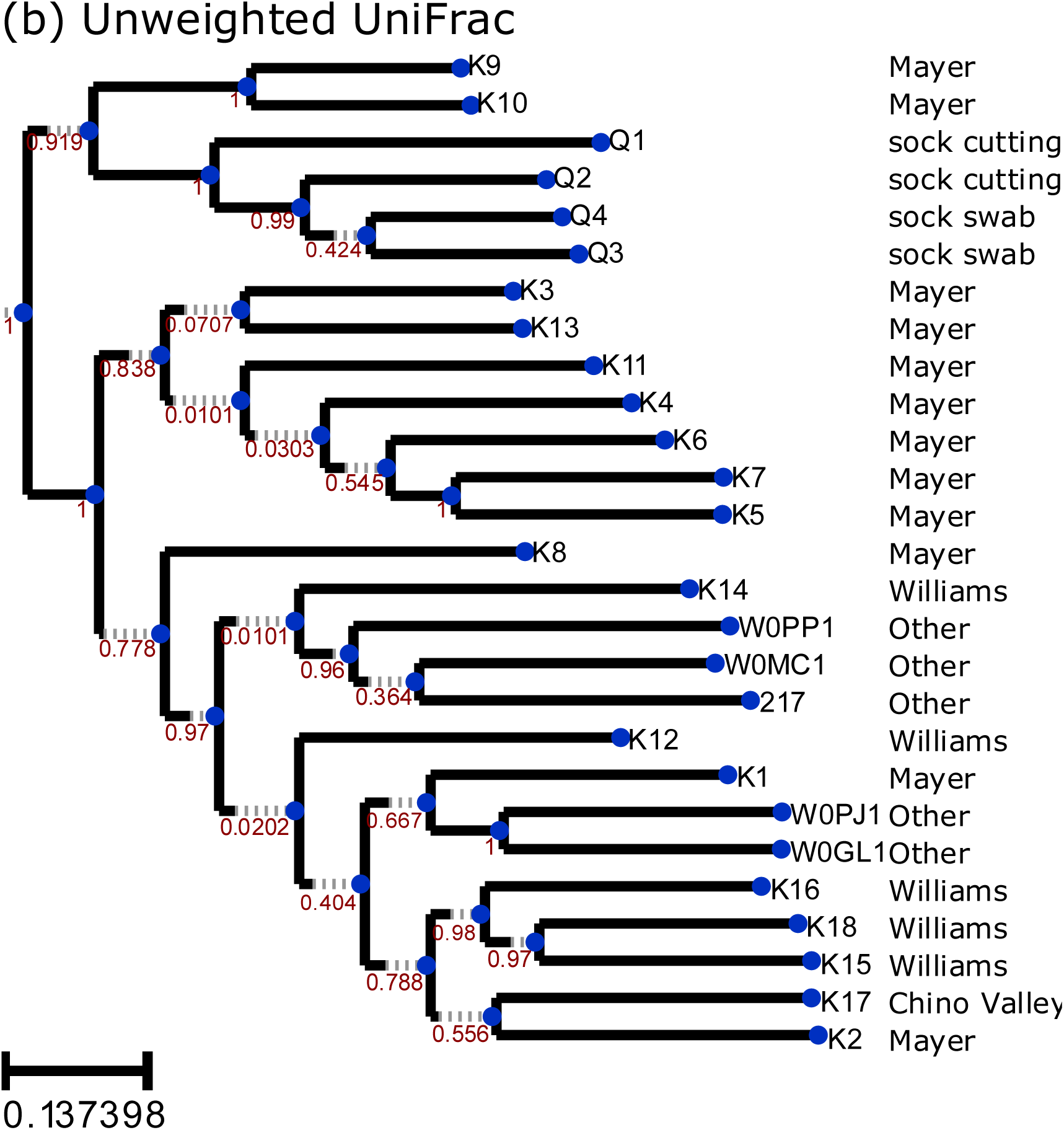
Neighbor joining trees illustrating (a) unweighted Unifrac and (b) weighted UniFrac distances between samples. Leaves of the trees represent samples, and the branch length between pairs of leaves represents the dissimilarity between samples. Values above the internal nodes of the tree represent jackknife support values, ranging between 0 and 1. Larger values indicate more robust groupings of samples.

Both the weighted and unweighted UniFrac neighbor joining trees illustrate that the sock samples are all most similar to each other in composition, and that the closest soil samples are all from Mayer, where the suspect left the remains of the victim. Clustering of the sock and Mayer soil samples was highly robust, and suggest that the soil on the suspect’s sock could have informed investigators of which cities should be the focus of search efforts.

Analysis of the taxonomic composition revealed typical soil microorganisms for all soil samples (Fig. 3). As would be expected, the dominant microorganisms in the sock sample were taxa commonly found on human skin. Because the victim’s body was left at sites K9 and K10, we were concerned that skin microbes found at those sites would link those samples to the sock, irrespective of the soil microbial composition. We therefore performed parallel analyses to those presented here after filtering the dominant skin bacterial family found here, Staphylococcaceae, from the sock and soil samples. This resulted in the taxonomic compositions presented in Fig. S2. The sock samples were still most similar to the Mayer samples, even after removal of all Staphylococcaceae (Fig. S1).

**FIG. 3.**
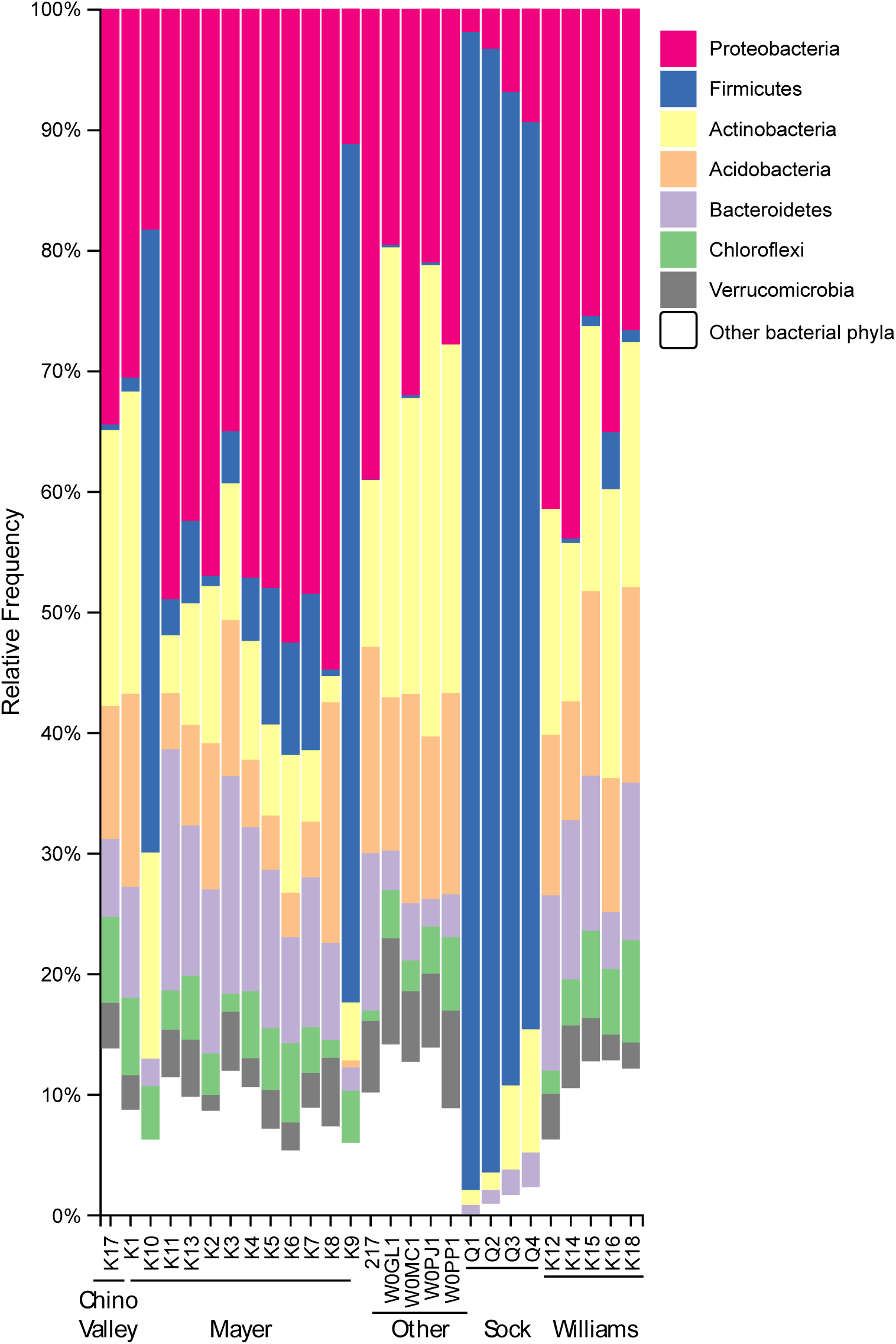
Microbiome taxonomic composition at the phylum levels for all samples.

The sock swabbing technique obtained more of the soil profile with less human-associated microbes compared to the cutting of the sock method (Fig. 3), though all of the sock samples clustered together in our analyses, suggesting that either approach would have led us to the same conclusion (Fig. 2). Because the swabbing technique produced less human-associated microbes and was not destructive of the evidence, this approach is likely a better choice than extraction of DNA from the sock cuttings.

Microbial DNA amplification was observed in the reagent blanks. This is to be expected as bacteria are ubiquitous in any environment so careful monitoring of contamination from laboratory equipment and reagents is crucial (27). Reagent blanks were processed alongside both the sock and reference soil samples. Although bacteria were present in both reagent blanks, the composition and abundance varied greatly from the reference soil samples and the query sock samples (Fig. S3).

## Discussion

The QIIME 2 platform was applied for analysis of microbiome data in this study. QIIME 2’s retrospective data provenance tracking feature may prove to be helpful in microbiome-based forensics work. All analysis steps, including versions of software installed on the system when each step was run, are automatically tracked as metadata associated with its results. This would allow an expert to determine with complete certainty what computational steps were taken to generate a result. As DNA analysis workflows can be complex, this automated recording will provide experts with the information they need to be confident in a given result or to identify potential issues such as the presence of a software bug or suboptimal analysis step in a workflow that may impact conclusions drawn from the data. Data provenance can be viewed for the results generated for this paper by loading the QIIME 2 results from Supplementary File 1 with QIIME 2 View (https://view.qiime2.org).

The ability to associate a piece of evidence to a location is a valuable tool to law enforcement. In the case presented here, a murder suspect was known to have traveled over a long distance during a few days’ time. Other evidence pointed to this individual having committed a murder, and both police and the victim’s family were anxious to discover the remains of the victim. With the police having narrowed down some possible locations for the body site, we were able to link an item of evidence to the location where the victim’s body was left. This result was possible because we were able to create a small database of locations known to have been visited by the suspect through police investigative techniques. Although the victim’s body was located with the assistance of the suspect, we believe that had this not happened, we would have been able to advise law enforcement that the soil embedded in the suspect’s sock most likely came from the Mayer, Arizona area rather than other locations where soil was collected based on the data presented here. In cases where areas coming in contact with the item of evidence are not known, a database of known soils from across a county or even a state would be very useful.

## Acknowledgements

Microbiome analysis and the QIIME 2 platform were partially funded by National Science Foundation grant number 1565100 to JGC. Laboratory supplies, sequencing, and personnel time were covered by the Arizona Board of Regents Technology Infrastructure Research Initiative Fund as startup funds to CMH.

Ecoss reference soil samples were collected with support by the National Science Foundation, Division of Environmental Biology (grant number: DEB-1241094).

## Supporting Information

**FIG. S1.**
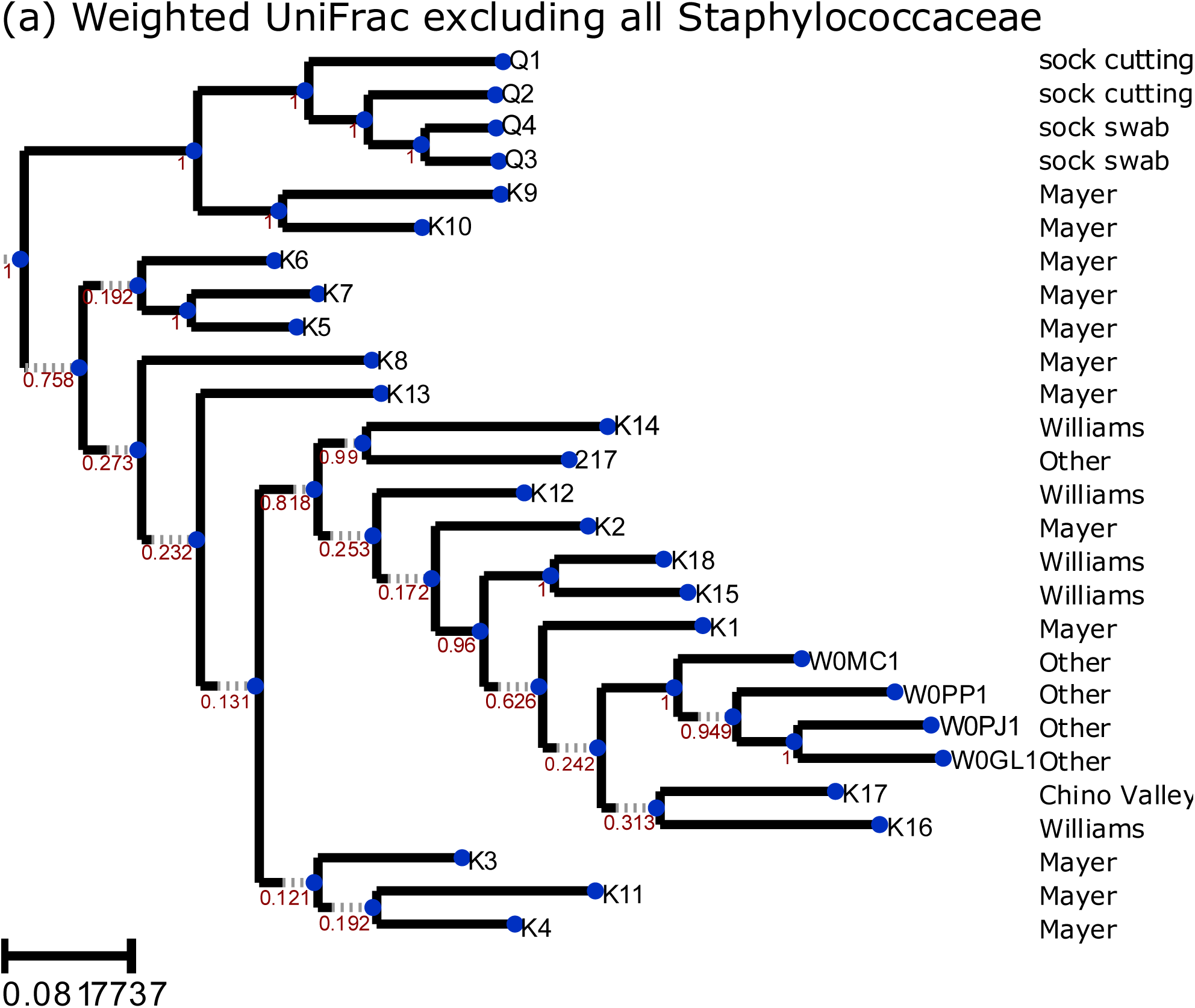
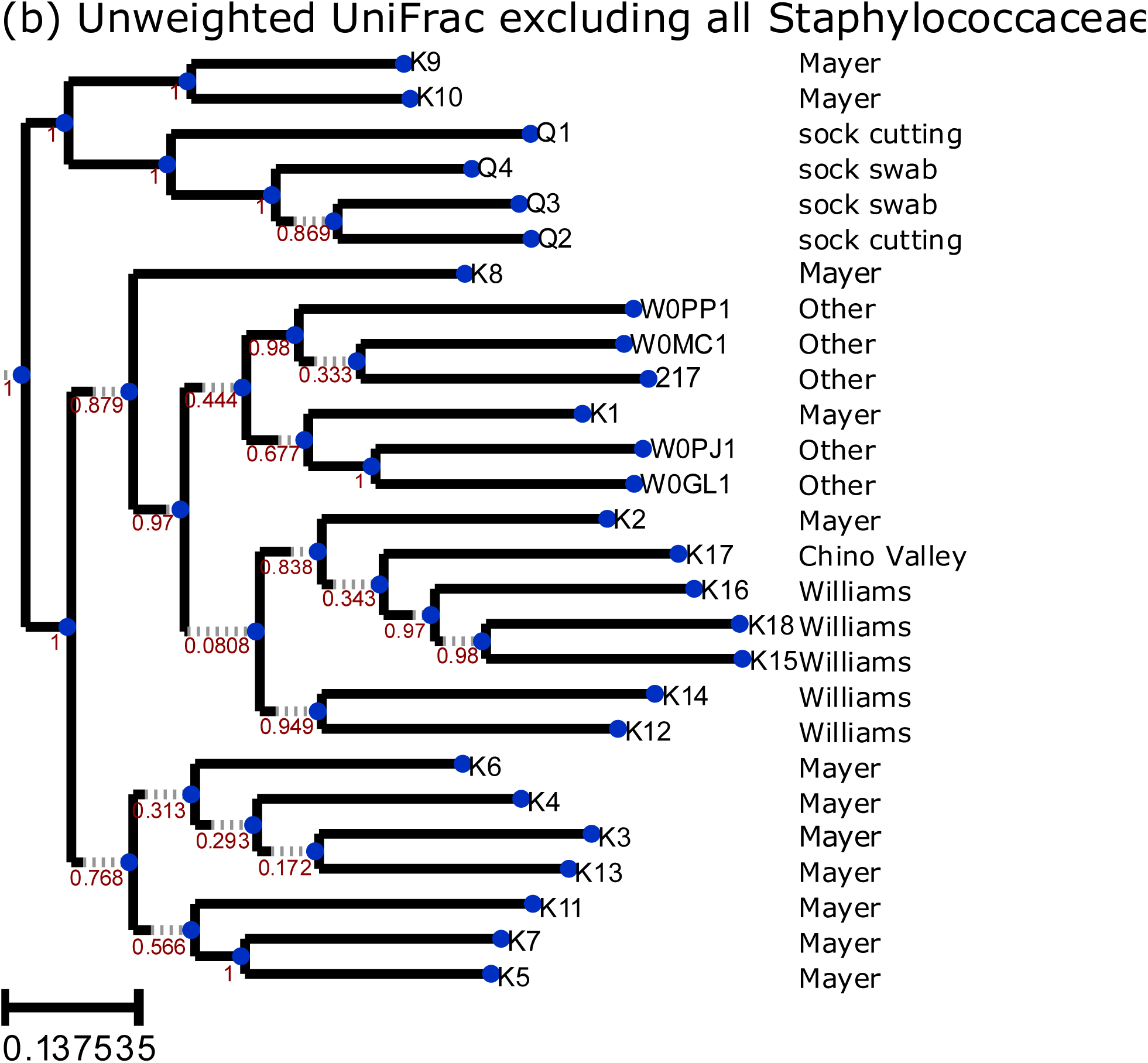
Neighbor joining trees illustrating (a) unweighted Unifrac and (b) weighted UniFrac distances between samples after excluding all Staphylococcaceae.

**FIG. S2.**
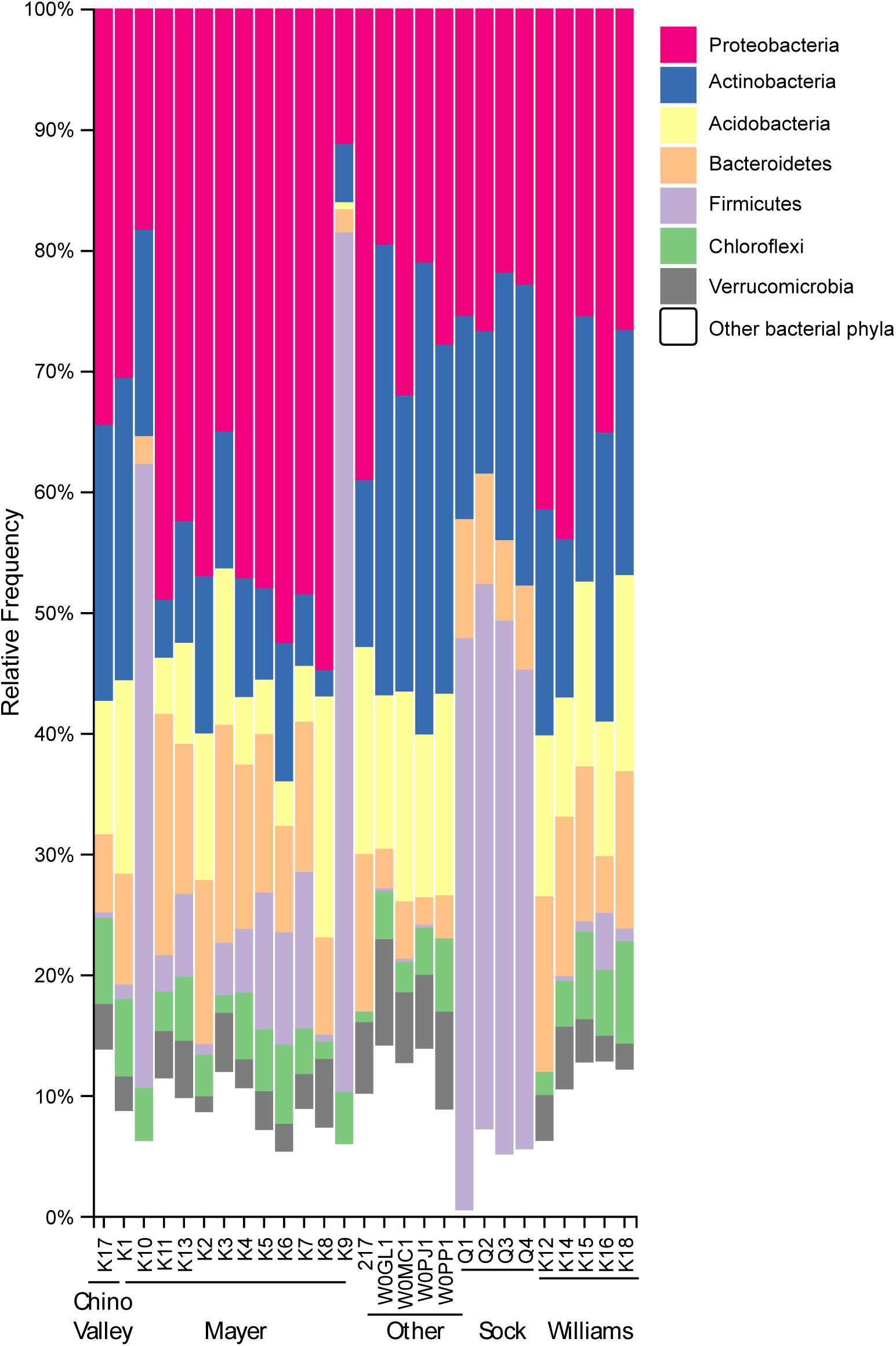
Microbiome taxonomic composition at the phylum levels for all samples after excluding all Staphylococcaceae.

**FIG. S3.**
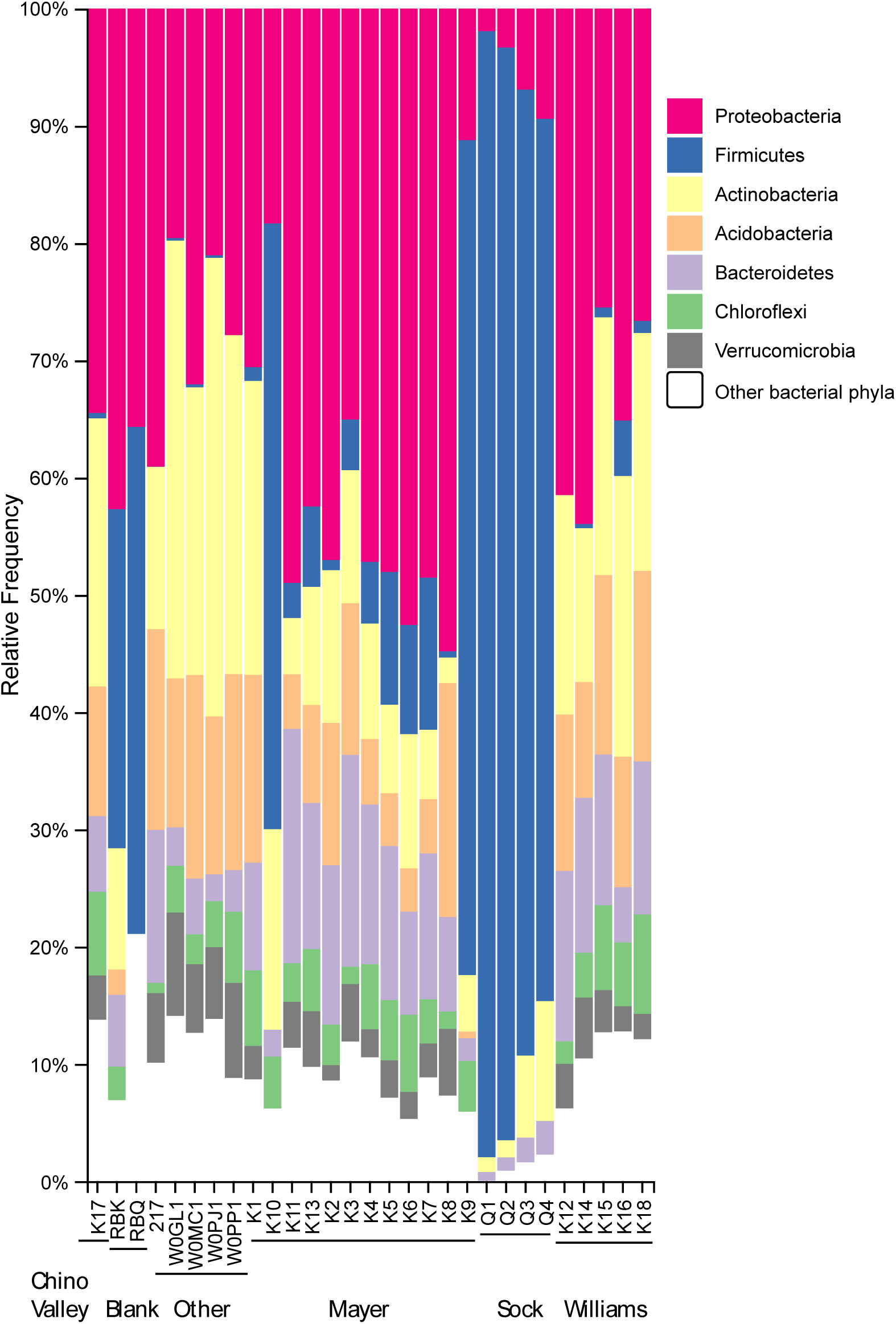
Microbiome taxonomic composition at the phylum levels for all samples including reagent blanks RBQ and RBK.

