## Supplementary File 1 for "Forensic analysis of Soil Microbiomes: Linking Evidence to a Geographic Location": draw-tree.ipynb

```
# %% [code]
import pandas as pd
from ete3 import Tree, TreeStyle, TextFace, faces
import os.path

# %% [markdown]
Load relevant metadata

# %% [code]
df = pd.read_csv('../../sample-metadata.tsv', sep='\t', index_col='sample-id')
df = df[df['project'] == 'flagstaff-pd']
df = df[df['Site'] != 'control']

# %% [code]
base_dir = '/Users/gregcaporaso/gdrive/manuscripts/2019.08-cocking-soil-forensics/'
fns = [
    ('wu-sample-clustering-nj.tre', '(a) Weighted UniFrac', 'figure-2a-unrooted.pdf'),
    ('uu-sample-clustering-nj.tre', '(b) Unweighted UniFrac', 'figure-2b-unrooted.pdf'),
    ('wu-no-staph-sample-clustering-nj.tre', '(a) Weighted UniFrac excluding all Staphylococcaceae', 'figure-s1a-unrooted.pdf'), 
    ('uu-no-staph-sample-clustering-nj.tre', '(b) Unweighted UniFrac excluding all Staphylococcaceae', 'figure-s1b-unrooted.pdf'),
]

for (input_fn, title, output_fn) in fns:
    input_fp = os.path.join(base_dir, input_fn)
    output_fp = os.path.join(base_dir, output_fn)
    t = Tree(newick=input_fp,
             format=0)
    ts = TreeStyle()
    ts.title.add_face(faces.TextFace(title, fsize=15), 0)
    ts.show_leaf_name = True
    ts.show_branch_support = True

    for sample_id in df.index:
        site = df['Site'][sample_id]
        t.get_leaves_by_name(sample_id)[0].add_face(TextFace(site), column=0, position = "aligned")


    t.render(output_fp, tree_style=ts)

# %% [code]
```
