## Supplementary File 1 for "Forensic analysis of Soil Microbiomes: Linking Evidence to a Geographic Location": pcoa-minus-staph.ipynb

```
# %% [code]
import qiime2
import qiime2.plugins.taxa
import qiime2.plugins.feature_table
import qiime2.plugins.diversity

import skbio

import pandas as pd

# %% [code]
sample_metadata = qiime2.Metadata.load('../../sample-metadata.tsv')
taxonomy = qiime2.Artifact.load('./taxonomy.qza')
table = qiime2.Artifact.load('./table.qza')
tree = qiime2.Artifact.load('rooted-tree.qza')

table = qiime2.plugins.feature_table.actions.filter_samples(table, metadata=sample_metadata, where="NOT [Site]='control'")[0]

# %% [code]
filtered_table = qiime2.plugins.taxa.actions.filter_table(table, taxonomy, exclude='f__Staphylococcaceae')[0]

# %% [code]
table.view(pd.DataFrame).shape

# %% [code]
filtered_table.view(pd.DataFrame).shape

# %% [code]
filtered_table.view(pd.DataFrame).sum(axis=1)

# %% [code]
rare_table = qiime2.plugins.feature_table.actions.rarefy(filtered_table, 1000)[0]

# %% [code]
dm = qiime2.plugins.diversity.actions.beta_phylogenetic(rare_table, tree, 'weighted_unifrac')[0]

# %% [code]
dm.view(skbio.DistanceMatrix)

# %% [code]
distances = dm.view(skbio.DistanceMatrix).to_data_frame()[['Q1', 'Q2', 'Q3', 'Q4']].sort_values(by='Q1')
distances['Site'] = sample_metadata.get_column('Site').to_series()
distances

# %% [code]
qiime2.plugins.diversity.visualizers.beta_rarefaction(table, 
                                                      metric='weighted_unifrac', 
                                                      clustering_method='nj', 
                                                      metadata=sample_metadata, 
                                                      sampling_depth=1000, 
                                                      phylogeny=tree)[0]

# %% [code]
qiime2.plugins.diversity.visualizers.beta_rarefaction(filtered_table, 
                                                      metric='unweighted_unifrac', 
                                                      clustering_method='nj', 
                                                      metadata=sample_metadata, 
                                                      sampling_depth=1000, 
                                                      phylogeny=tree)[0]


# %% [code]
```
