## Supplementary File 1 for "Forensic analysis of Soil Microbiomes: Linking Evidence to a Geographic Location": view-distance-vector.ipynb

```
# %% [code]
import qiime2

# %% [code]
weighted_unifrac_dm = qiime2.Artifact.load('./core-metrics-results/weighted_unifrac_distance_matrix.qza')

# %% [code]
weighted_unifrac_dm

# %% [code]
import skbio

# %% [code]
dm = weighted_unifrac_dm.view(skbio.DistanceMatrix)

# %% [code]
dm

# %% [code]
dm.to_data_frame()['Q4'].sort_values()

# %% [code]
```
