## Supplementary File 1 for "Forensic analysis of Soil Microbiomes: Linking Evidence to a Geographic Location": README.txt

The .qza and .qzv files in this directory can be viewed using QIIME 2 View (https://view.qiime2.org). To see the provenance of any of those files, select the "Provenance" tab after opening a .qza or .qzv file on QIIME 2 View.
